# Sequence-context dependency shown by eIF5^G31R^ mutant for UUG start codon selection in *Saccharomyces cerevisiae*

**DOI:** 10.1101/2023.02.09.527861

**Authors:** Anup Kumar Ram

## Abstract

In higher eukaryotes, the efficiency of start codon selection depends upon its sequence context A/Gxx**<u>AUG</u>**G, especially at the −3 and +4 position. However, *S. cerevisiae* prefers AAA/U**<u>AUG</u>**U sequence context surrounding the AUG start codon. Furthermore, it was demonstrated that the first AUG codon on the mRNA serves as a signal for ribosomal recognition of the initiation site selection. In yeast, the G31R mutation in eIF5 shows Suppressor of initiation codon (Sui^−^) phenotype enabling 40S ribosome to initiate at UUG start codon. To understand whether the UUG codon recognition by the eIF5^G31R^ mutant is sensitive to the sequence context, we made different *HIS4-LacZ* reporter constructs with UUG as start codon and changed the sequence context at −3 and −2 positions. The HIS4^UUG^-LacZ transcripts that carry purine (A/G) at the −3-position showed better ß-Galactosidase activity than pyrimidine (U/C). Furthermore, on changing the −1 position for the AAA context, we discovered purines are more preferred at the −1 position for the UUG start codon selection.

## 1 Introduction

In eukaryotes, the sequence context surrounding the AUG start codon profoundly impacts the translation initiation efficacy. Marilyn Kozak showed that the AUG start codon that harbors purine at −3 and guanine residue at +4 position (A/Gxx<u>AUG</u>G) is most efficiently translated in the eukaryotes (Kozak 1986; Kozak 1991; Kozak 1991). The importance of −3 and +4 positions was experimentally tested using pre-pro-insulin model system and shown that a point mutation at either of these positions causes decrease in the proinsulin expression levels. The introduction of C/G/A at −3 nucleotide (nt) position and keeping A at +4 nt position causes drastic reduction in the proinsulin translation efficiency. A similar decrease in proinsulin levels were observed when the C/U nucleotide was introduced at the −3 nt position with a constant of G at +4 nt position. These observations suggest that a single base change can affect translational initiation efficiency (Kozak 1984; Kozak 1986). The sequence context surrounding the AUG start codon in the *S. cerevisiae* varies slightly from the Kozak consensus sequence and reportedly uses AAA/U<u>AUG</u>U sequence context for the efficient AUG start codon selection (Hamilton et al. 1987; Cigan and Donahue 1987). As the AUG start codon occupies the P-site of the 40S ribosome, the −3 to −1 upstream sequence resides in the E-site of the 40S ribosome, where it interacts with the eIF2α subunit, small ribosomal proteins Rps5 and Rps26 (Sharifulin et al. 2013; Visweswaraiah et al. 2015; Visweswaraiah and Hinnebusch 2017). Evolutionarily, most of the AUG start codons reside in a good or optimal context owing to its expression. However, the eIF1 mRNA transcript carries an AUG start codon in a sub-optimal context (CGT<u>ATG</u>T), which negatively regulates its own translation when the eIF1 concentration is above a certain threshold (Martin-Marcos et al. 2014; Benitez-Cantos et al. 2020). Similarly, the expression of the eIF1 and eIF5 in mammalian systems is auto-regulated by the context surrounding the AUG start codon (Loughran et al. 2012; Li et al. 2017). Translation initiation from the non-AUG codons in eukaryotes are rare. However, prokaryotes are known to initiate translation from the non-AUG codons; for example, an analysis of 620 bacterial genomes revealed that 8% of the mRNAs initiates translation using UUG as the start codon and 12% mRNA initiates translation using GUG start codon (Villegas and Kropinski 2008; Cao et al. 2021). In yeast, translation of the mitochondrial version of glycyl tRNA synthetase (*GRS1*) commences at the UUG start codon (Chen et al. 2008). The sequence context of the UUG start codon in the *GRS1* mRNA plays an important role in its efficient translation, where the presence of AAA nucleotide at −3 to −1 position is critical (Chen et al. 2008). However, there are no reports on whether the recognition of the non-AUG codon for translation initiation by the Sui^−^ mutants is contingent on the sequence context. It is intriguing to investigate whether the UUG codon recognition by the eIF5^G31R^ mutant is dependent on the sequence context surrounding the UUG start codon.

In yeast, eIF5 plays an important role in translation start site selection through its GAP (GTPase activating protein) function and by sustaining eIF1 in the 40s initiation complex avoiding Pi to come out until and unless there is proper AUG codon-anti codon interaction. G31R mutation in eIF5, makes it Sui^−^ mutant causing preferential utilization of UUG as initiation codon (D. and T. 1993; Donahue and Cigan 1988; Saini et al. 2014). The eIF5^G31R^ and eIF2β^S264Y^ Sui^−^ mutants did not initiate at CUG or GUG near-cognate start codons, however, they prefer initiation at the UUG near cognate start codon (Antony A and Alone 2018). But till now, there are no reports on whether the recognition of the non-AUG codon for translation initiation by the Sui^−^ mutants is contingent on the sequence context. It is intriguing to investigate whether the UUG codon recognition by the eIF5^G31R^ mutant is dependent on the sequence context surrounding the UUG start codon.

## 2. Materials and Methods

### 2.1. Strains

The *Saccharomyces cerevisiae* yeast strain used in this study was: YP823; *Mat α, ura3-52, leu2-3,112, trp1Δ63, GCN2*+ (Antony A and Alone 2017).

### 2.2. Plasmid and Oligonucleotides used in this study

Plasmids and oligonucleotides used in this study are listed in Table 1 and Table 2 respectively.

**Table 1.** Plasmids used in this study

| Sr. No. | Plasmid number | Plasmid name | Type | Reference |
| --- | --- | --- | --- | --- |
| 1 | A823 | pYcplac22 | Single copy<br>(s.c) | (Gietz et al. 1988) |
| 2 | A838 | pYcplac22-eIF5<sup>G31R</sup> | s.c | (Ram et al. 2022) |
| 3 | A1361 | pYcplac33-HIS4-AAA_AUG_LacZ | s.c | (Ram et al. 2022) |
| 4 | A1362 | pYcplac33-HIS4-UUU_AUG_LacZ | s.c | (Ram et al. 2022) |
| 5 | A1363 | pYcplac33-HIS4-AAA_UUG_LacZ | s.c | (Ram et al. 2022) |
| 6 | A1364 | pYcplac33-HIS4-UUU_AUG_LacZ | s.c | (Ram et al. 2022) |
| 7 | A1365 | pYcplac33-HIS4-CUU_UUG_LacZ | s.c | This study |
| 8 | A1366 | pYcplac33-HIS4-CGU_UUG_LacZ | s.c | This study |
| 9 | A1367 | pYcplac33-HIS4-CCU_UUG_LacZ | s.c | This study |
| 10 | A1368 | pYcplac33-HIS4-GUU_UUG_LacZ | s.c | This study |
| 11 | A1369 | pYcplac33-HIS4-GAU_UUG_LacZ | s.c | This study |
| 12 | A1370 | pYcplac33-HIS4-GGU_UUG_LacZ | s.c | This study |
| 13 | A1371 | pYcplac33-HIS4-GCU_UUG_LacZ | s.c | This study |
| 14 | A1372 | pYcplac33-HIS4-AUU_UUG_LacZ | s.c | This study |
| 15 | A1373 | pYcplac33-HIS4-AGU_UUG_LacZ | s.c | This study |
| 16 | A1374 | pYcplac33-HIS4-UAU_UUG_LacZ | s.c | This study |
| 17 | A1375 | pYcplac33-HIS4-UGU_UUG_LacZ | s.c | This study |
| 18 | A1376 | pYcplac33-HIS4-UCU_UUG_LacZ | s.c | This study |
| 19 | A1378 | pYcplac33-HIS4-CAU_UUG_LacZ | s.c | This study |
| 20 | A1379 | pYcplac33-HIS4-AAU_UUG_LacZ | s.c | This study |
| 21 | A1380 | pYcplac33-HIS4-ACU_UUG_LacZ | s.c | This study |
| 22 | A1436 | pYcplac33-HIS4-AAU_UUG_LacZ | s.c | This study |
| 23 | A1437 | pYcplac33-HIS4-AAU_UUG_LacZ | s.c | This study |
| 24 | A1438 | pYcplac33-HIS4-AAU_UUG_LacZ | s.c | This study |

**Table 2.**
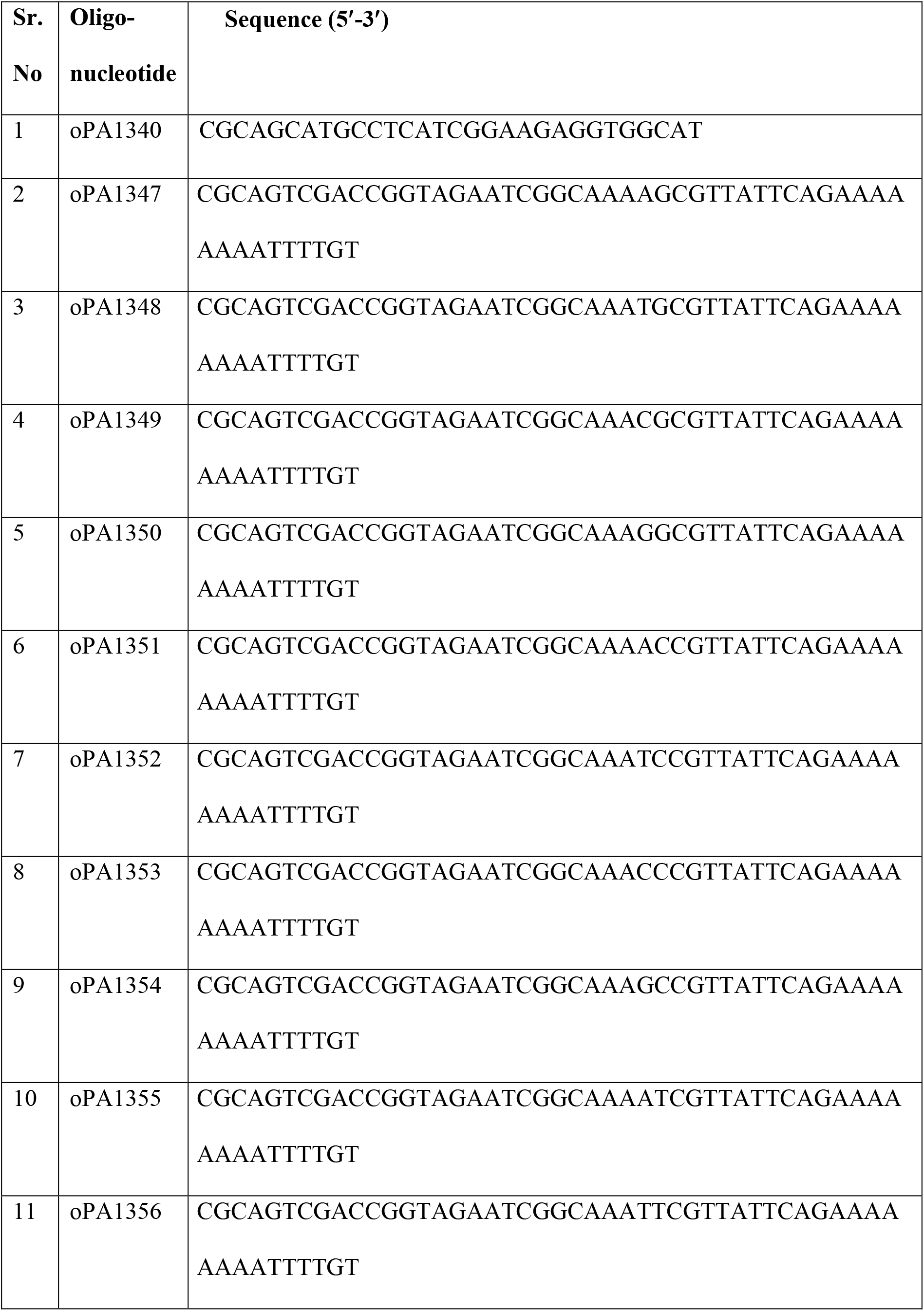
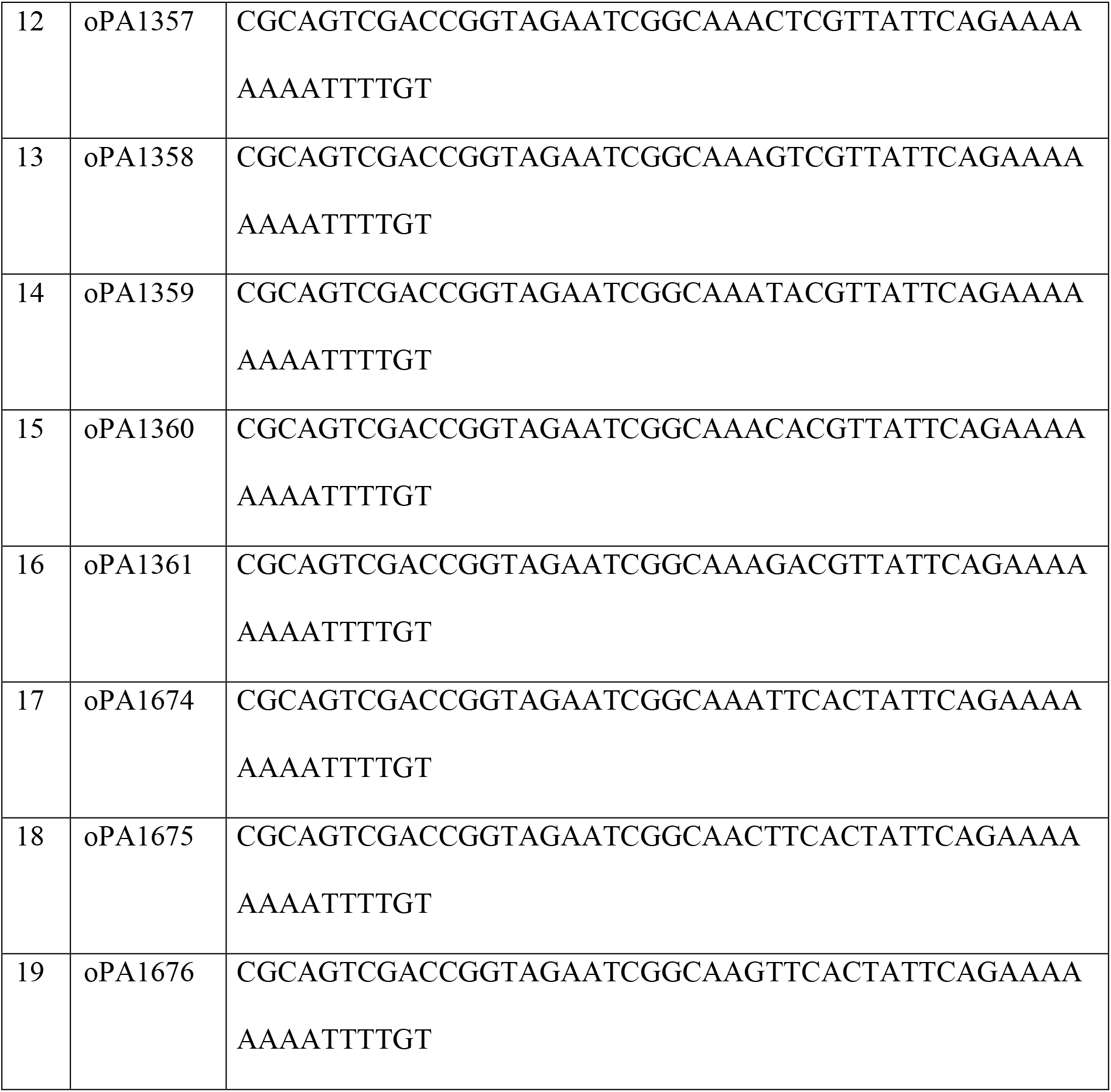
Oligonucelotides used in this study

### 2.3. Construct

The pYcplac33-His4-lacZ reporter construct was generated as follows- the *LacZ* region and *HIS4* 3’ UTR region were PCR amplified using oligonucleotides combination oPA1009/oPA1010 and oPA1341/oPA1342 respectively and cloned at SalI-BamHI and BamHI-KpnI sites. The 541 bp of *HIS4* promoter and the N-terminal 7 amino acids region was PCR amplified using oligonucleotide combination oPA1340 with oPA1347/oPA1348/oPA1349/oPA1350/oPA1351/oPA1352/oPA1353/oPA1354/oPA1355/oPA1356/oPA1357/oPA1358/oPA1359/oPA1360/oPA1361/oPA1674/oPA1675/oPA1676 cloned into vector pYcplac33(A309) at HindIII-SalI site for CUU/CAU/CGU/CCU/GUU/GAU/GGU/GCU/AUU/AAU/AGU/ACU/UAU/UGU/UCU/AAU/AAG/AAC sequence context respectively having UUG as start codon.

### 2.4. β-Galactosidase reporter assay

Yeast cells were transformed with appropriate reporter plasmids. Five colonies from each transformant were grown overnight at 30°C with shaking at 220 rpm in synthetic dextrose (SD) media with appropriate nutrient supplements and grown till OD_600_ ~ 0.8. Cells were harvested by centrifugation at 13,000xg for 20 minutes at 4°C and resuspended in LacZ buffer (60 mM Na_2_HPO_4_, 40 mM NaH_2_PO_4_, 10 mM KCl, and 1 mM MgSO_4_, pH 7.0) and cells were mechanically lysed by vortexing using acid-washed glass beads (200 μm) at 4 °C. Clarified cell extract (~30 μg) was mixed with LacZ buffer, followed by addition of 180 μl of ONPG (4 mg/ml in LacZ buffer). After 30 minutes of incubation, absorbance was measured at 420 nm (Molecular Devices Spectra Max). Protein estimation was done using Bradford assay and the β-galactosidase activity (nmol of O-nitrophenyl-β-D-galactopyranoside cleaved per min per mg) was calculated after normalization with total protein.

## 3. Results

### 3.1. Selection of the UUG start codon by the eIF5^G31R^ mutation is sequence context-dependent

Our earlier work established that the eIF5^G31R^ mutation shows a strong Sui^−^ phenotype (Antony A and Alone 2018). Next, we asked if the sequence context surrounding the UUG start codon plays any significant role in the UUG start codon selection in the eIF5^G31R^ mutant. Previously, we made HIS4-LacZ reporter constructs having start codon either AUG or UUG with change in sequence context from −3 to −1 position, where AAA constitutes good, and UUU constitutes a poor sequence context. As per HIS4-LacZ reporter data, the eIF5^G31R^ mutation showed a ~2.6-fold decrease in UUG codon recognition efficacy when the UUG codon was in the poor sequence context (Ram et al. 2022).

To further delineate the effect of the different nucleotides at −3 and −2 positions on the efficacy of UUG start codon recognition by the eIF5^G31R^ mutant, we made 15 different possible HIS4-LacZ reporter constructs. These 15 *HIS4-LacZ* constructs were transformed into yeast strain (YP823) carrying either empty vector or eIF5^G31R^ mutation construct. The constructs having purine at the −3 position showed higher *HIS4-LacZ* reporter activity compared to the constructs having pyrimidine at the −3 position (Fig. 1A & 1B). With either A or G at the −3 position there were four possible constructs, out of those four constructs 3/4^th^ times A is being favored (as AAU, ACU, and AUU sequence context) and 2/4^th^ times G is being favored (as GAU and GCU sequence context) at −3 position for UUG start codon selection by eIF5^G31R^ mutant as represented by the bit map (Fig. 2). Among these five favored sequence contexts, four contain A or C at the −2 position, which is also observed for the AUG and UUG start codon selection in yeast as reported previously (Li et al 2017; Chen et al 2008). However, in the other sequence context, GGU, GUU and AGU did not show any significant increase in the β- galactosidase reporter activity even after having purine at the −3 position. It is likely that the G or U nucleotide at the −2 position is not compatible with the A/G nucleotide at the −3 position for the UUG start codon selection. The UUG codon recognition in GCU context was almost 2/3 of AAA context, suggesting G at −3 and C at −2 nucleotide position is the second best after AA at −3/-2 nucleotide position for UUG start codon selection in eIF5^G31R^ mutant.

**Fig. 1.**
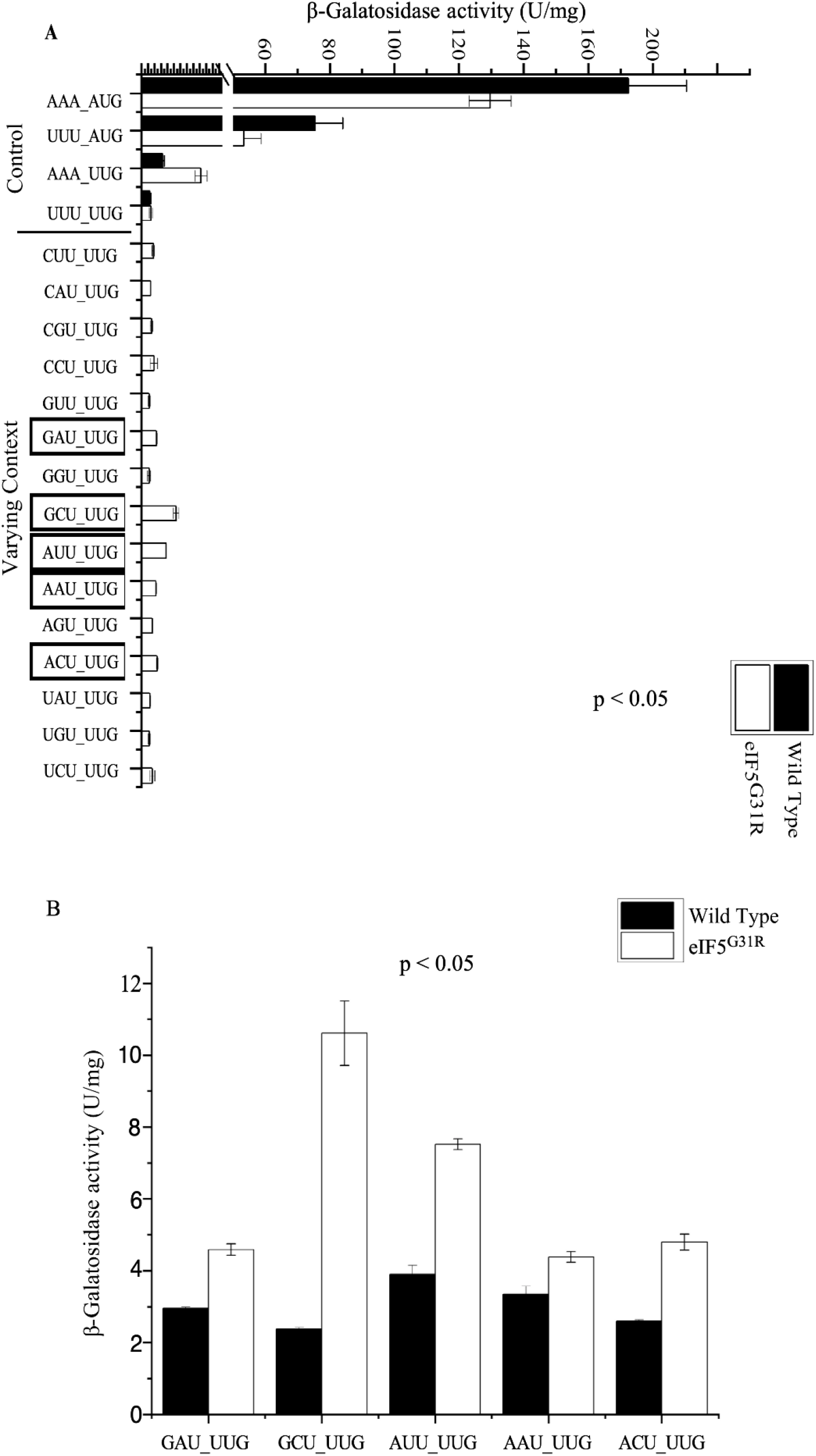
Sequence context governs UUG codon recognition by eIF5^G31R^ mutant. (A)Yeast strain YP823 carrying either empty vector pYcplac22 (A823) & pYcplac22-eIF5^G31R^ (A838) construct was transformed with HIS4^AUG^-LacZ good/poor context (A1361/62), HIS4^UUG^-LacZ good/poor context (A1363/64) as experimental control. YP823 having pYcplac22-eIF5^G31R^ was further transformed with 15 other reporter constructs having different nucleotide combination (as varying context) at the −3 and −2 nucleotide position (A1365/66/67/68/69/70/71/72/73/74/75/76/78/79/80). All these transformants were grown till OD600 ~ 0.8 in SD plus leucine medium at 30°C. WCE were prepared and β-galactosidase activity (nmol of O-nitrophenyl-b-D-galactopyranoside cleaved per min per mg) was measured and plotted as bar graph. (B) Yeast strain YP823 carrying either empty vector (pA823) & SUI5 (pA838) construct was transformed with five selected HIS4^UUG^-LacZ constructs (A1369/71/72/79/80), and the β-galactosidase assay was performed.

**Fig. 2.**
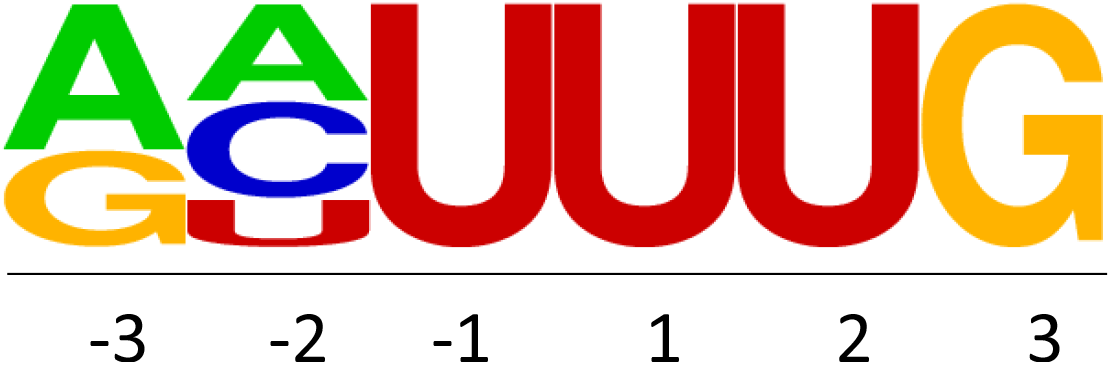
The favored sequence context at −3 and −2 nucleotide for UUG codon recognition by eIF5^G31R^ mutant.

### 3.2. Sui5 favours purine at −1 position for UUG start codon selection

To test the importance of −1 nucleotide position in the UUG codon selection in the eIF5^G31R^ mutant, we made three more *HIS4-LacZ* reporter constructs where nucleotide sequence at the −3 to −1 is either AAG, AAU, or AAC. These *HIS4-LacZ* reporter constructs were transformed into the YP823 yeast strain along with either empty vector or eIF5^G31R^ mutation. The *HIS4-LacZ* data suggests that the preference for nucleotide at the −1 position is of the following order A > G > U > C for selecting the UUG start codon in the eIF5^G31R^ mutant compared with the WT control. This data suggests that the eIF5^G31R^ mutant strongly favors AAA nucleotide at the −3 to −1 position for the UUG start codon selection (Fig. 3).

**Fig. 3.**
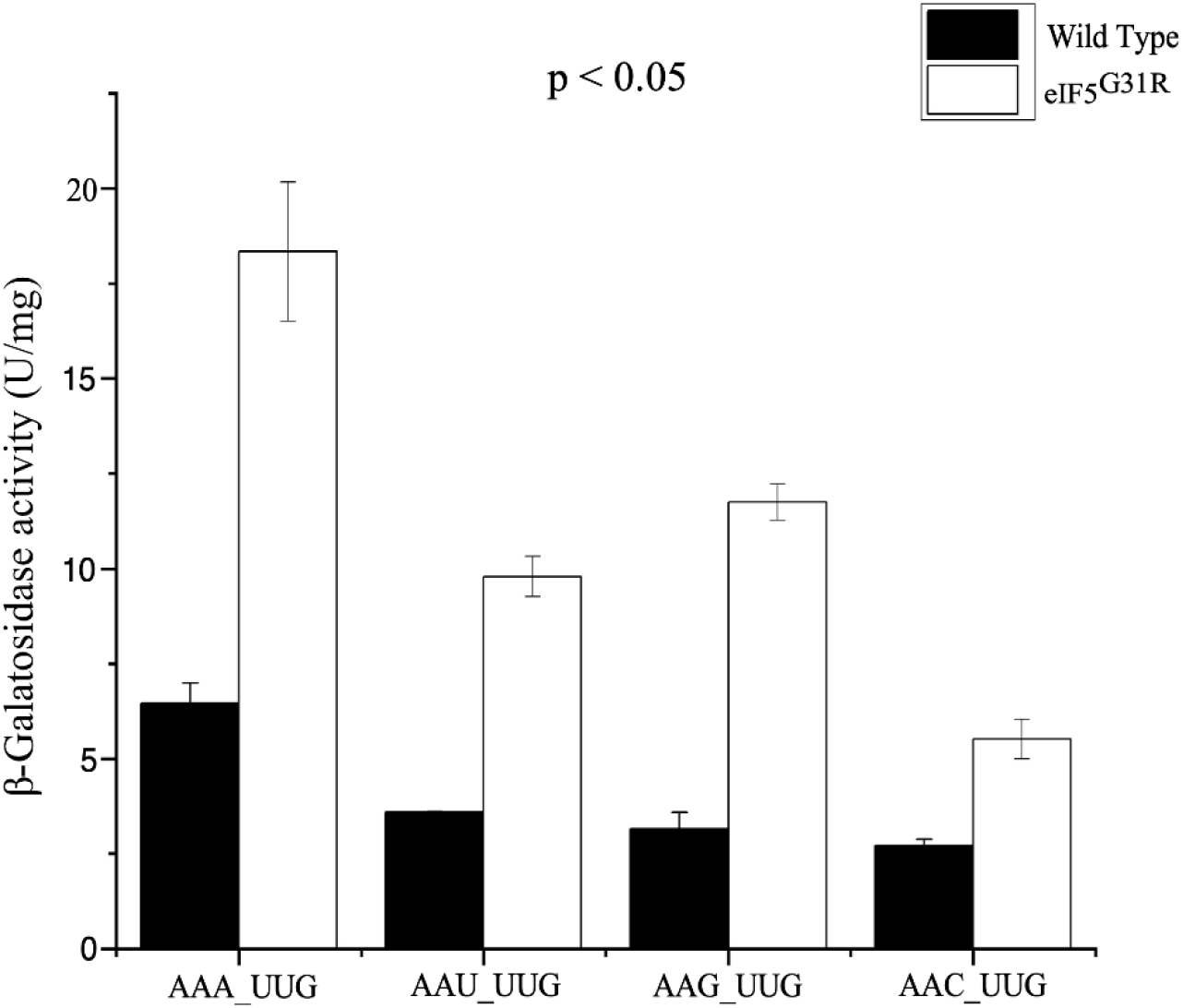
The nucleotide at the −1 position regulates UUG codon recognition by eIF5^G31R^ mutant. Yeast strain YP823 carrying either empty vector (pA823) & SUI5 (pA838) construct was transformed with derivatives of HIS4^UUG^-LacZ construct having a change in −1 nucleotide position (A1361, A1436, A1437 & A1438), and the β-galactosidase assay was performed.

## 4. Discussion

Previous work reported that the sequence context surrounding the AUG start codon plays a significant role in the start codon selection in the higher eukaryotes compared to yeast (Hinnebusch 2011; Thakur et al. 2020). However, translation initiation using the UUG codon is facilitated by the sequence context in the *GRS1* transcript (Chen et al. 2008). Our data suggest that the eIF5^G31R^ mutant has a strong penchant for initiating translation from the UUG codon compared to the CUG or GUG codon. We observed that the sequence context surrounding the UUG start codon is critical for recognition by the eIF5^G31R^ mutant. The UUU sequence context at the −3 to −1 position has a negative effect on UUG codon recognition by the eIF5^G31R^ mutant. In contrast, the AAA sequence context shows maximum initiation from the UUG start codon by the eIF5^G31R^ mutant. Individual nucleotides substitution at −3 to −1 position reveals preference of purines at these locations. We conclude that the UUG start codon selection by the eIF5^G31R^ mutant is sequence context-dependent. Genome-wide ribosomal profiling of the eIF5^G31R^ mutant may give us more clarity about the most used context by the initiating ribosome at the UUG codon.

## Acknowledgments

I thank Dr. Pankaj V Alone, NISER Bhubaneswar for his guidance and helpful discussion all throughout this work.

## Author contributions

AKR has designed and performed the experiments. AKR has analyzed the data, written the manuscript and arranged the figures.

